# SARS-CoV-2 spike protein-mediated cell signaling in lung vascular cells

**DOI:** 10.1101/2020.10.12.335083

**Authors:** Yuichiro J. Suzuki, Sofia I. Nikolaienko, Vyacheslav A. Dibrova, Yulia V. Dibrova, Volodymyr M. Vasylyk, Mykhailo Y. Novikov, Nataliia V. Shults, Sergiy G. Gychka

**Affiliations:** Department of Pharmacology and Physiology, Georgetown University Medical Center, Washington, DC 20007 USA; Department of Pathological Anatomy N2, Bogomolets National Medical University, Kiev, 01601 Ukraine; Centralized Department of Pathological Anatomy of Ivano-Frankivsk District Clinical Hospital, Ivano-Frankivsk, 76000 Ukraine; The Regional Municipal Institution «Sumy District Forensic Medical Examination Bureau», Sumy, 40050 Ukraine

**Keywords:** cell signaling, coronavirus, COVID-19, SARS-CoV-2, vascular

## Abstract

Currently, the world is suffering from the pandemic of coronavirus disease 2019 (COVID-19), caused by severe acute respiratory syndrome coronavirus 2 (SARS-CoV-2) that uses angiotensin-converting enzyme 2 (ACE2) as a receptor to enter the host cells. So far, 30 million people have been infected with SARS-CoV-2, and nearly 1 million people have died because of COVID-19 worldwide, causing serious health, economical, and sociological problems. However, the mechanism of the effect of SARS-CoV-2 on human host cells has not been defined. The present study reports that the SARS-CoV-2 spike protein alone without the rest of the viral components is sufficient to elicit cell signaling in lung vascular cells. The treatment of human pulmonary artery smooth muscle cells or human pulmonary artery endothelial cells with recombinant SARS-CoV-2 spike protein S1 subunit (Val16 – Gln690) at 10 ng/ml (0.13 nM) caused an activation of MEK phosphorylation. The activation kinetics was transient with a peak at 10 min. The recombinant protein that contains only the ACE2 receptor-binding domain of SARS-CoV-2 spike protein S1 subunit (Arg319 – Phe541), on the other hand, did not cause this activation. Consistent with the activation of cell growth signaling in lung vascular cells by SARS-CoV-2 spike protein, pulmonary vascular walls were found to be thickened in COVID-19 patients. Thus, SARS-CoV-2 spike protein-mediated cell growth signaling may participate in adverse cardiovascular/pulmonary outcomes, and this mechanism may provide new therapeutic targets to combat COVID-19.

## 1. Introduction

Coronaviruses are positive sense single stranded RNA viruses that often cause the common cold [1,2]. Some coronaviruses can, however, be lethal. Currently, the world is suffering from the pandemic of coronavirus disease 2019 (COVID-19) caused by severe acute respiratory syndrome coronavirus 2 (SARS-CoV-2) [3,4]. So far, 30 million people have been infected with SARS-CoV-2 worldwide, causing serious health, economical, and sociological problems. SARS-CoV-2 uses angiotensin converting enzyme 2 (ACE2) as a receptor to enter the host cells [5,6]. Lung cells are the primary targets of SARS-CoV-2, resulting in severe pneumonia and acute respiratory distress syndrome (ARDS) [7,8]. So far, nearly 1 million people have died because of COVID-19.

It has been noted that certain populations of infected individuals are severely affected by and die of COVID-19. Elderly patients with systemic hypertension and other cardiovascular diseases are particularly susceptible to developing severe and possibly fatal conditions [4,9,10]. Thus, managing the pulmonary and cardiovascular aspects of COVID-19 is considered to be the key for reducing the severity of COVID-19 and the associated mortality. However, it is unclear exactly how SARS-CoV-2 affects humans. Thus, understanding the mechanism of SARS-CoV-2 actions should help develop therapeutic strategies to reduce the mortality and morbidity associated with COVID-19.

The SARS-CoV-2 spike protein is critical to initiate the interactions between the virus and the host cell surface receptor and facilitates the viral entry into the host cell by assisting in the fusion of the viral and host cell membranes [11,12]. This protein consists of two subunits: Subunit 1 (S1) that contains the ACE2 receptor-binding domain (RBD) and Subunit 2 (S2) that is responsible for fusion.

The current dogma of the SARS-CoV-2 mechanism is that the binding of viral spike protein to the host ACE2 receptor results in the entry of virus into the host cells and the cellular response is a result of the viral infection [5,6,11,12]. However, we herein show that SARS-CoV-2 spike protein alone without the rest of the viral components is sufficient to elicit cell signaling in human host cells, suggesting a novel biological mechanism of SARS-CoV-2 actions.

## 2. Materials and Methods

### 2.1 Cell culture

Human pulmonary artery smooth muscle cells and human pulmonary artery endothelial cells were purchased from ScienCell Research Laboratories (Carlsbad, CA, USA), and rat pulmonary artery smooth muscle cells were purchased from Cell Applications (San Diego, CA, USA). Cells were cultured in accordance with the manufacturers’ instructions in 5% CO_2_ at 37°C. Cells at passages 3–6 were maintained in low fetal bovine serum (0.4%)-containing medium overnight before the treatment as routinely performed in experiments on cell signaling and protein phosphorylation [13].

Cells were treated with the recombinant SARS-CoV-2 spike protein full length S1 subunit that contains most of the S1 subunit (Val16 - Gln690) with a molecular weight of Ū75 kDa (RayBiotech, Peachtree Corners, GA, USA) or the recombinant SARS-CoV-2 spike protein RBD (RayBiotech) that only contains the RBD region (Arg319 - Phe541) with a molecular weight of Ū25 kDa. Some cells were pretreated for 1 h with the rabbit anti-ACE2 antibody (Catalog # 4355; Cell Signaling Technology, Danvers, MA).

### 2.2 Western blotting

To prepare cell lysates, cells were washed in phosphate buffered saline and solubilized with lysis buffer containing 50 mM Hepes (pH 7.4), 1% (v/v) Triton X-100, 4 mM EDTA, 1 mM sodium fluoride, 0.1 mM sodium orthovanadate, 1 mM tetrasodium pyrophosphate, 2 mM PMSF, 10 μg/ml leupeptin, and 10 μg/ml aprotinin. Samples were then centrifuged at 16,000*g* for 10 min at 4°C, supernatants collected, and protein concentrations determined [13].

Equal amounts of protein samples were electrophoresed through a reducing sodium dodecyl sulfate polyacrylamide gel. Proteins were then electro-transferred to the Immobilon-FL Transfer Membrane (MilliporeSigma, Burlington, MA, USA). The membrane was blocked with Odyssey blocking buffer (LI-COR, Lincoln, NE, USA) for one hour at 25°C and incubated overnight with the rabbit anti-phospho-MEK1/2 (Ser217/221) antibody (Catalog #9154; Cell Signaling), the mouse anti-MEK1/2 antibody (Catalog #4694; Cell Signaling), the rabbit anti-phospho-Akt (Ser473) antibody (Catalog #9271; Cell Signaling), or the rabbit anti-phospho-Stat3 (Tyr705) antibody (Catalog #9131; Cell Signaling) at 4°C. Washed membranes were then incubated with IRDye 680RD or IRDye 800CW (LI-COR) for one hour. Signals were obtained by using the Odyssey Infrared Imaging System (LI-COR).

### 2.3 Histological examinations of patient samples

Postmortem lung tissues were collected from 10 patients who died of COVID-19 in Ukraine during the period of March - July, 2020. For comparison, we used the archival materials of lung tissues from 10 patients who died of influenza A (H1N1) during the epidemic in November and December of 2009. Clinical studies were approved by the regional committee for medical research ethics in Kiev, Ukraine and performed under the Helsinki Declaration of 1975 revised in 2013 or comparable ethical standards.

For histological examinations, samples of autopsy materials approximately 10 mm thick were fixed overnight in 10% buffered formalin at room temperature. Fixed tissues were embedded in paraffin. From paraffin blocks, 5 μm thick sections were made using a microtome. The sections were stained with hematoxylin and eosin. Histological specimens were examined using a Leica BX 51 microscope, a Leica MC 190 digital camera, and the Leica LAS software at magnifications of 100x - 200x. Morphometric investigations included the assessment of pulmonary arterial wall thickness and pulmonary arterial lumen index (the ratio of internal vessel area to external vessel area) using the ImageJ software. Seven vessels analyzed for each patient.

### 2.4 Statistical Analysis

For Western blotting data, means and standard errors of mean (SEM) were computed. Two groups were compared by a two-tailed Student’s *t* test, and differences between more than two groups were determined by the analysis of variance (ANOVA). *p* < 0.05 was defined to be statistically significant. For the morphometric analysis, IBM SPSS Statistics software version 22.0 was used for the statistical calculations. Mann-Whitney U Test was used to define the statistical significance at *p* < 0.05.

## 3. Results

### 3.1 SARS-CoV-2 spike protein alone can activate cell signaling in human cells

To test the hypothesis that SARS-CoV-2 spike protein alone can elicit cell signaling, human pulmonary artery smooth muscle cells were treated with the full length S1 subunit (Val16 - Gln690) of the SARS-CoV-2 spike protein for 0, 10, and 30 min. As shown in Fig. 1A, the spike protein at a concentration of 10 ng/ml (0.133 nM) strongly activated the phosphorylation of 45 kDa MEK at Ser217 and Ser221 residues. The kinetics of MEK phosphorylation promoted by the full-length S1 subunit (Val16 - Gln690) of the SARS-CoV-2 spike protein was consistently found to be transient with a peak at 10 min. This fast activation suggests that this may be a receptor-mediated cell-signaling event. Similarly, full-length S1 subunit SARS-CoV-2 spike protein promoted the phosphorylation of MEK in human pulmonary artery endothelial cells (Fig. 1B). SARS-CoV-2 spike protein, however, did not activate other signaling events such as Akt (Fig. 1A) and Stat3 (Fig. 1B) pathways.

**Fig. 1:**
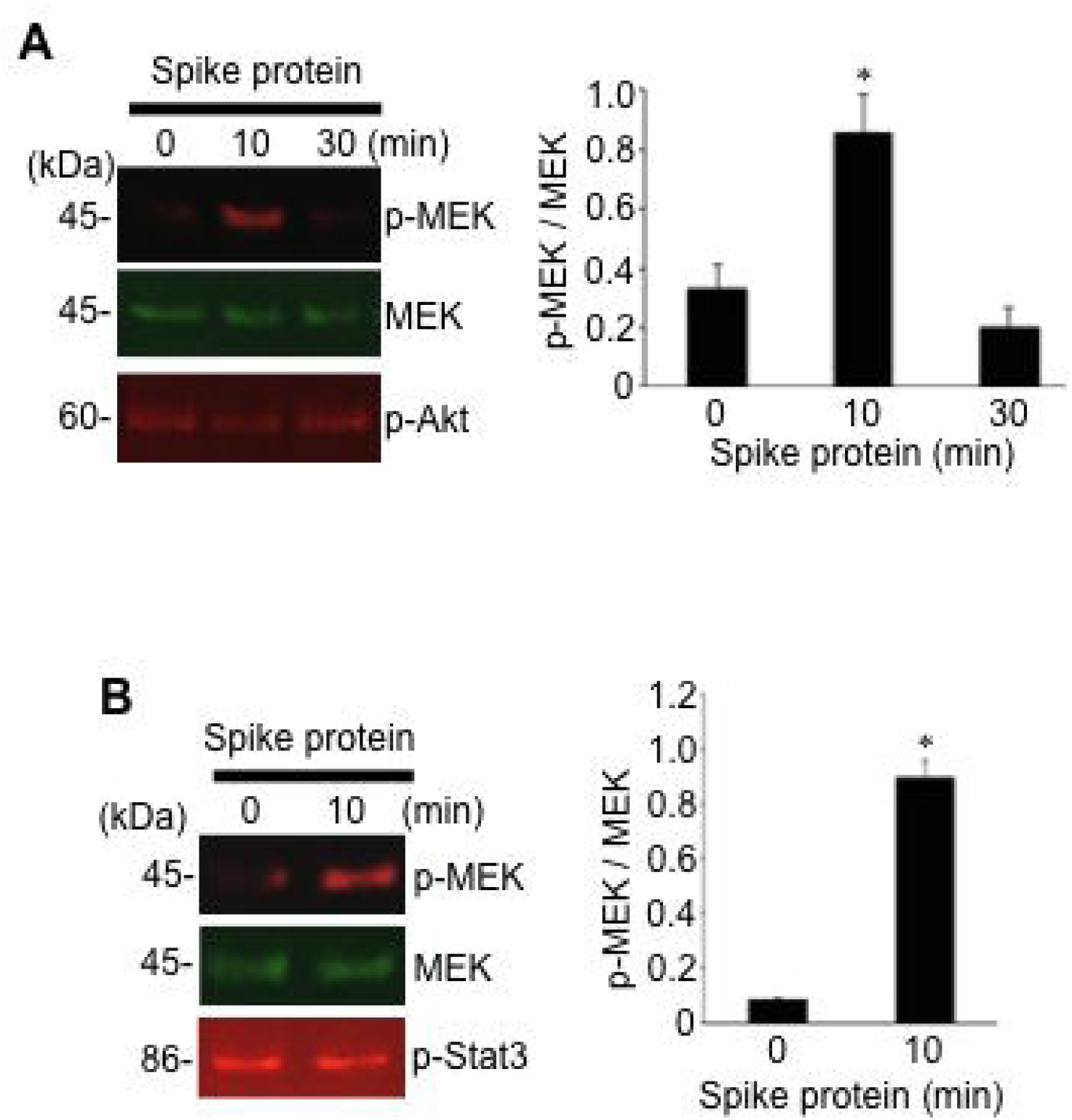
SARS-CoV-2 spike protein S1 promotes the MEK phosphorylation in human cells. (A) Human pulmonary artery smooth cells (N=5) and (B) human pulmonary artery endothelial cells (N=3) were treated with recombinant full length S1 subunit of SARS-CoV-2 spike protein (Val16 - Gln690) at 10 ng/ml for durations indicated. Cell lysates were prepared and subjected to Western blotting using antibodies against phosphorylated MEK (p-MEK), MEK protein, phosphorylated Akt (p-Akt) and phosphorylated Stat3 (p-Stat3). Bar graphs represent means ± SEM. *Significantly different from 0 min at *p* < 0.05.

As we performed experiments to determine the effects of the full-length S1 subunit of SARS-CoV-2 spike protein on rat pulmonary artery smooth muscle cells, we surprisingly found that, not only SARS-CoV-2 spike protein did not promote the MEK phosphorylation, but rather decreased the phosphorylation. As shown in Fig. 2, the treatment of full length SARS-CoV-2 spike protein S1 resulted in the dephosphorylation of MEK as early as 10 min after the treatment and this dephosphorylation event was maintained for at least 60 min. Thus, SARS-CoV-2 spike protein promotes the MEK phosphorylation in human cells, but not in rat cells.

**Fig. 2:**
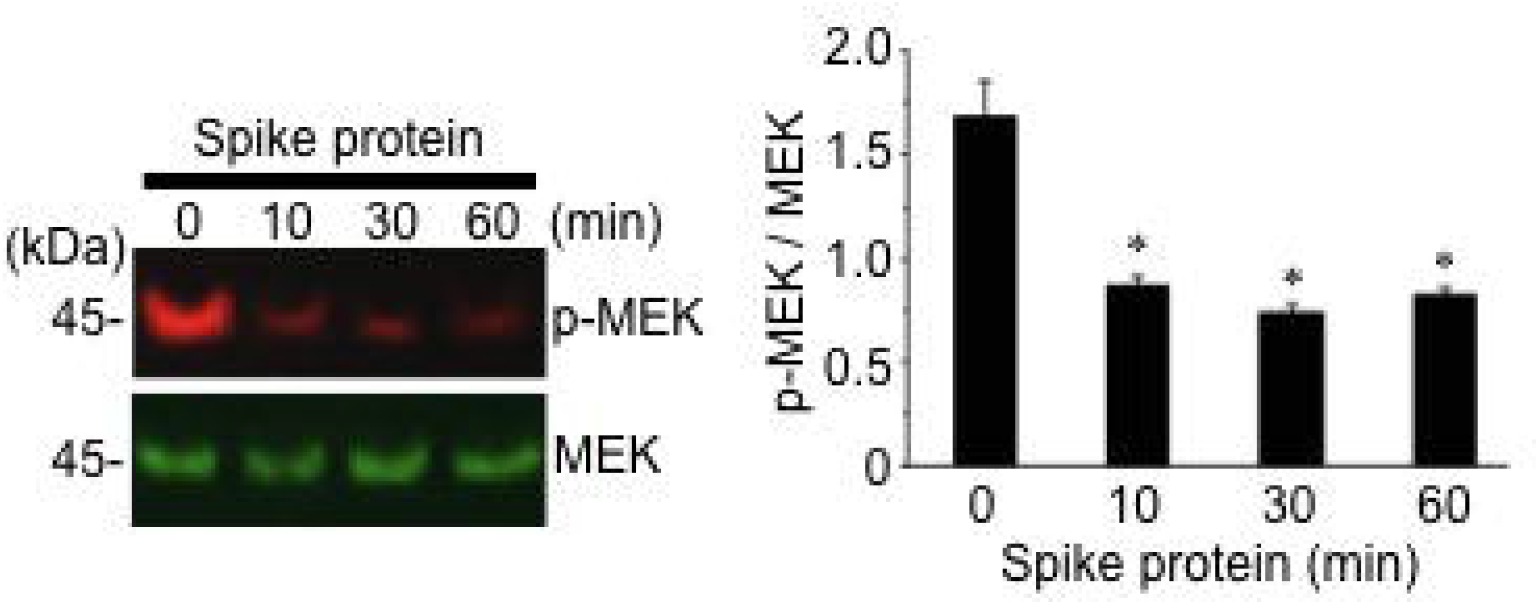
In rat cells, SARS-CoV-2 spike protein S1 does not phosphorylate, rather dephosphorylates MEK. Rat pulmonary artery smooth muscle cells were treated with recombinant full length S1 subunit of SARS-CoV-2 spike protein (Val16 - Gln690) at 10 ng/ml for durations indicated. Cell lysates were prepared and subjected to Western blotting using antibodies against phosphorylated MEK (p-MEK) and MEK protein. The bar graph represents means ± SEM (N = 4). *Significantly different from 0 min atp < 0.05.

To confirm that the action of SARS-CoV-2 spike protein is through its well-known receptor ACE2,^5,6^ human pulmonary artery smooth muscle cells were pretreated with the ACE2 antibody for 1 hour before treating with the full-length S1 subunit of SARS-CoV-2 spike protein. The ACE2 antibody alone caused the activation of MEK, and SARS-CoV-2 spike protein did not further increase this MEK phosphorylation signal (Fig. 3).

**Fig. 3:**
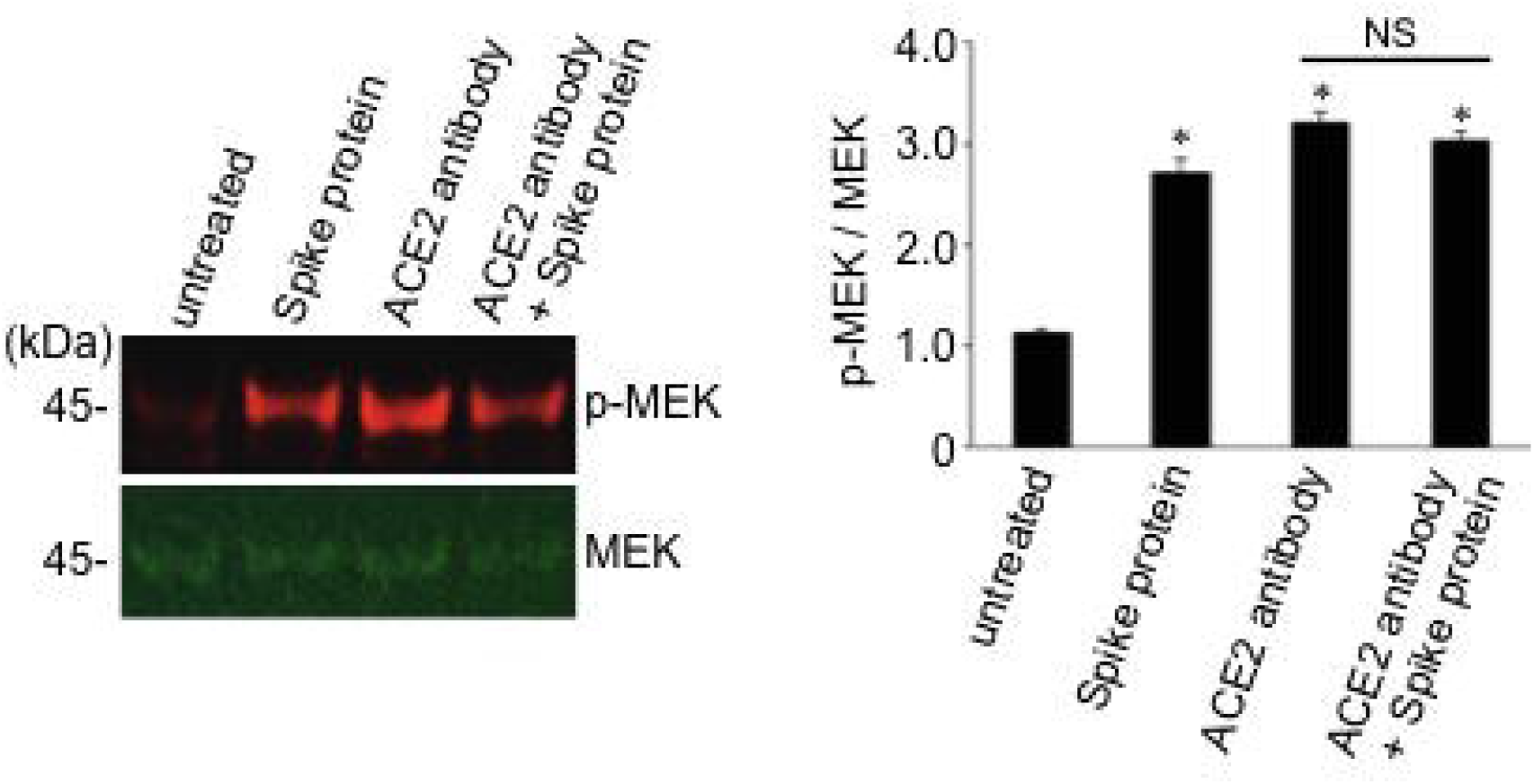
SARS-CoV-2 spike protein S1 does not promote the phosphorylation of MEK in the presence of the neutralizing antibody against ACE2. Human pulmonary artery smooth muscle cells were pre-treated with the ACE2 antibody for 1 hour and then treated with recombinant full length S1 subunit of SARS-CoV-2 spike protein (Val16 - Gln690) for 10 min. Cell lysates were prepared and subjected to Western blotting using antibodies against phosphorylated MEK (p-MEK) and MEK protein. The bar graph represents means ± SEM (N = 4). *Significantly different from untreated control at *p* < 0.05. NS denotes that the two values are not significantly different from each other at *p* < 0.05.

### 3.2 RBD domain alone is not sufficient to activate cell signaling

To test whether the RBD binding to ACE2 is sufficient to stimulate cell signaling for MEK phosphorylation, human cells were treated with the recombinant S1 RBD of SARS-CoV-2 spike protein that only contains the RBD (Arg319 - Phe541). In contrast to the full-length S1 subunit (Val16 - Gln690) that strongly phosphorylated MEK, S1 RBD (Arg319 - Phe541) did not activate the MEK phosphorylation in human pulmonary artery smooth muscle cells (Fig. 4A) or in human pulmonary artery endothelial cells (Fig. 4B). Thus, other regions of the spike protein in addition to RBD may be required for eliciting cell signaling for MEK phosphorylation.

**Fig. 4:**
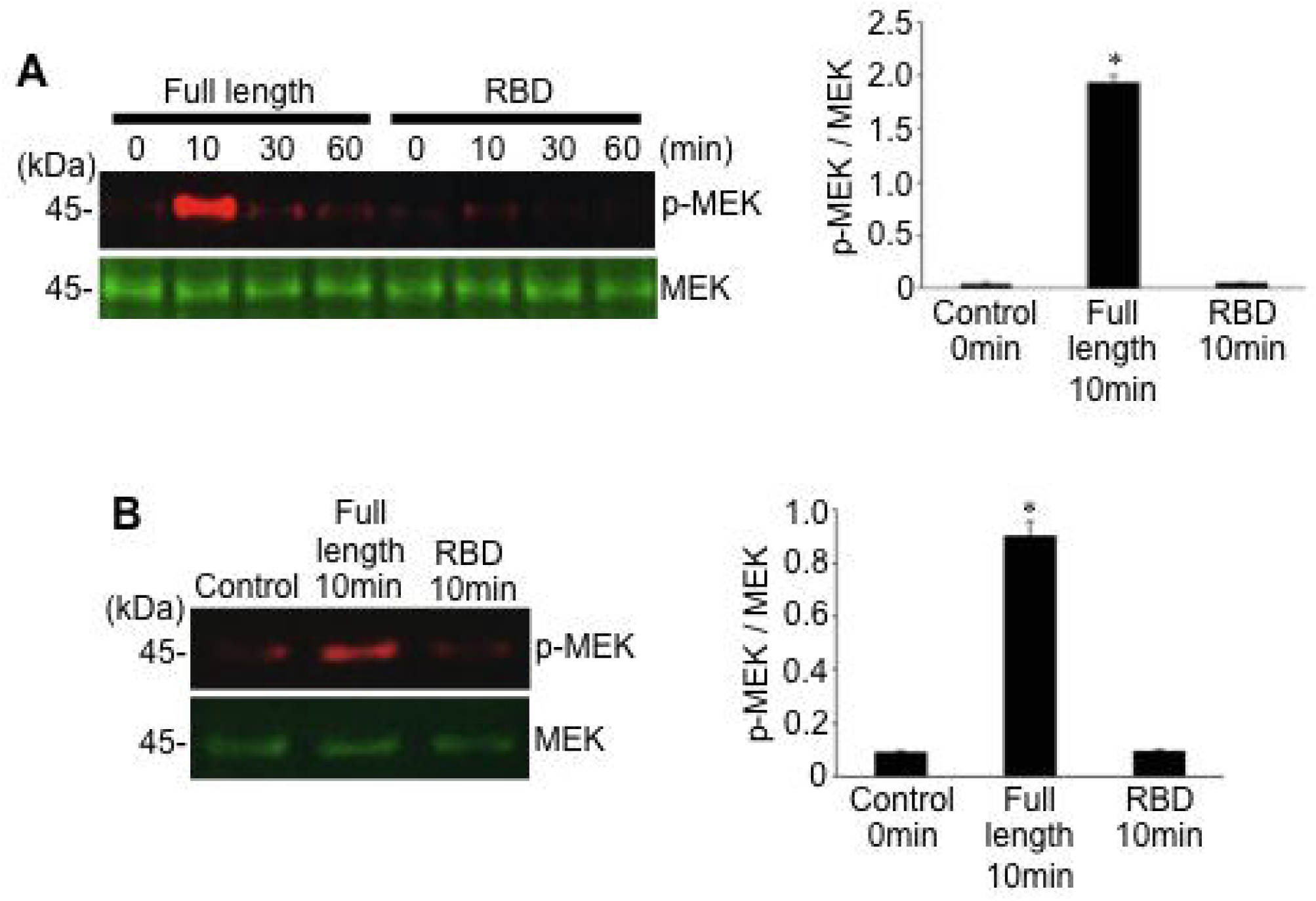
RBD only-containing SARS-CoV-2 spike protein S1 does not activate the MEK phosphorylation. (A) Human pulmonary artery smooth muscle cells and (B) human pulmonary artery endothelial cells were treated with recombinant full length S1 subunit (Val16 - Gln690) or the RBD region of S1 subunit (Arg319 - Phe541) at 100 ng/ml for durations indicated. Cell lysates were prepared and subjected to Western blotting using antibodies against phosphorylated MEK (p-MEK) and the MEK protein. Bar graphs represent means ± SEM (N = 3). *Significantly different from 0 min control at *p* < 0.05.

### 3.3 Pulmonary vascular walls are thickened in COVID-19 patients

Our results showing that SARS-CoV-2 spike protein is capable of activating the MEK/ERK pathway in pulmonary artery smooth muscle and endothelial cells suggest that cell growth signaling may be triggered in the pulmonary vascular walls in response to SARS-CoV-2. To test this, we examined the lung histology results of patients who died of COVID-19.

Fig. 5A shows the representative pulmonary vessels of patients who died of COVID-19. These pulmonary arteries consistently exhibit histological characteristics of vascular wall thickening. The walls of the pulmonary arteries are thickened mainly due to the hypertrophy of the tunica media. The vessels have lost the clarity of the boundaries with the surrounding lung parenchyma. The enlargement of smooth muscle cells of the middle lining of the arteries is observed. The nuclei of smooth muscle cells also are swollen, increased in size, have become rounded, and vacuoles are identified in the cytoplasm.

**Fig. 5:**
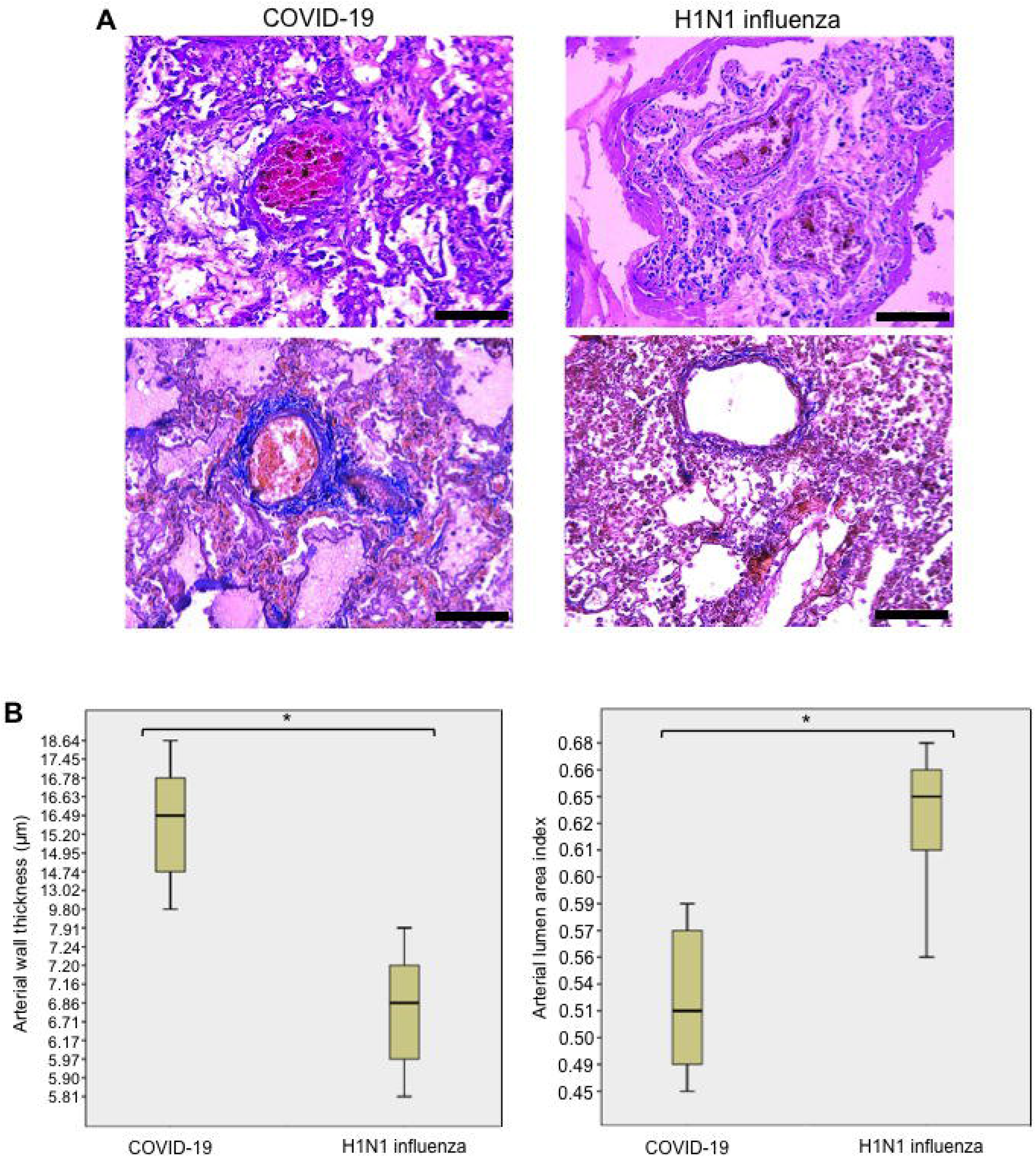
Pulmonary arterial walls are thickened in COVID-19 patients. (A) Representative H&E staining images of pulmonary arteries of patients who died of acute respiratory distress syndrome due to COVID-19 and H1N1 influenza. In COVID-19 patients, but not in H1N1 influenza patients, thickening of the pulmonary arterial walls was observed. Scale bars indicate 100 μm. (B) Box plots represent quantifications of pulmonary arterial wall thicknesses and lumen areas in COVID-19 and H1N1 influenza patients using H&E stained slides. The number of patients used for these analyses was 10 for COVID-19 and 10 for influenza. *Mann-Whitney U Test indicated that the two values are significantly different at p = 0.001.

By contrast, pulmonary vessels of patients who died of H1N1 influenza infection showed that there were no occurrences of thickened pulmonary arteries (Fig. 5A). Unlike COVID 19, smooth muscle cells form a compact layer of tunica media. There are no signs of the enlargement of smooth muscle cells.

We performed a morphometric analysis of the pulmonary vessels of patients who died of SARS-CoV-2 and H1N1 influenza virus. The median wall thickness value of the pulmonary arteries of COVID-19 patients was calculated to be 15.4 μm, while that of patients with H1N1 influenza was 6.7 μm (Fig. 5B, left), indicating that pulmonary arterial walls are thicker in COVID-19 patients more than 2-fold compared to influenza patients. These two values are significantly different from each other with *p*= 0.001. The pulmonary arterial lumen area was found to be significantly smaller in COVID-19 patients compared to influenza patients with *p*= 0.001 (Fig. 5B, right).

## 4. Discussion

The major finding of this study is that the SARS-CoV-2 spike protein without the rest of the virus can elicit cell signaling, specifically the activation of the MEK/ERK pathway, in human host lung vascular smooth muscle and endothelial cells. MEK is a mitogen-activated protein kinase kinase (MAPKK) that phosphorylates and activates extracellular-regulated kinase (ERK), one type of mitogen-activated protein kinases (MAPK). In this MEK/ERK pathway of cell signaling, MEK is activated by the phosphorylation by Raf1 kinase [14]. The MEK/ERK pathway is a well-known cell growth mechanism [15] and has also been shown to facilitate the viral replication cycles [16].

By contrast to the full-length S1 subunit of SARS-CoV-2 spike protein (Val16 - Gln690) that strongly activated MEK, RBD only containing protein (Arg319 - Phe541) did not activate MEK in human pulmonary artery smooth muscle or in endothelial cells. These results suggest that the protein regions other than the RBD (i.e. Val16 - Arg319 and/or Phe541 - Gln690) are required for eliciting cell signaling for the MEK phosphorylation. Thus, we propose that SARS-CoV-2 spike protein-mediated cell signaling is not merely cells responding to anything binding to the membrane surface protein, but is a growth factor/hormone-like specific cell signal transduction event that is well coordinated by the RBD as well as other protein regions that facilitate cell signaling. Our ACE2 antibody experiments as described in Fig. 3 support the involvement of ACE2. However, the binding of the ACE2 antibody to ACE2 seems sufficient to cause the activation of MEK. While we interpreted that the ACE2 antibody blocked and interfered with the binding of the RBD to ACE2, it is possible that ACE2 antibody-mediated cell signaling may have desensitized the subsequent spike protein-mediated effects to activate MEK. Thus, SARS-CoV-2 spike protein-mediated cell signaling may occur through a receptor other than ACE2.

In contrast to the effects of SARS-CoV-2 spike protein on human cells to activate the MEK phosphorylation, this protein promoted the dephosphorylation of MEK in rat cells. This species difference in the SARS-CoV-2 spike protein actions may be important to understand how this virus severely affects humans. We previously reported a novel ligand-mediated dephosphorylation mechanism of MEK induced by neurotensin and neuromedin N [17]. The SARS-CoV-2 spike protein signaling in rat pulmonary artery smooth muscle cells represents another example of the MEK dephosphorylation mechanism. These results also suggest that the species-specific actions of SARS-CoV-2 may depend on how spike protein-mediated cell signaling mechanisms differ among species.

It has been noted that elderly patients with systemic hypertension and other cardiovascular diseases are particularly susceptible to developing severe and possibly fatal conditions of COVID-19 [4,9,10]. Thus, the pathology of COVID-19 does not seem to be explained merely by the action of SARS-CoV-2 to enter the host cells for destruction. The present study shows that cell growth signaling may be triggered by this virus in both cultured cells and in COVID-19 patient samples. We propose that SARS-CoV-2 spike protein-mediated cell signaling promotes the hyperplasia and/or hypertrophy of vascular smooth muscle and endothelial cells, contributing to the complex cardiovascular outcomes in COVID-19. Further studies investigating this concept are warranted in order to develop effective therapeutic strategies to reduce the mortality associated with COVID-19.

Our histology results showing that COVID-19 patients have thickened pulmonary arterial walls also have clinical significance. These changes may reflect the tissue responses of the lungs to hypoxia and may play a vital role in the development of acute respiratory failure in COVID-19 patients. These results also indicate the possibility that patients who recovered from COVID-19 may be predisposed to developing pulmonary arterial hypertension and right-sided heart failure. We also noticed that published lung histology images of patients who died of ARDS during the 2002 - 2004 SARS outbreak due to the infection with SARS-CoV-1 did not exhibit the signs of thickened pulmonary vascular walls [18,19]. Thus, pulmonary vascular wall thickening is a unique feature of the SARS-CoV-2 infection and COVID-19.

## 5. Conclusions

Our results in human pulmonary artery smooth muscle and endothelial cells revealed that the SARS-CoV-2 spike protein S1 subunit is sufficient to trigger biological responses in the human host cells in the absence of the participation of the rest of the SARS-CoV-2 viral particle. We also found that the pulmonary vascular walls are thickened in COVID-19 patients. We propose that SARS-CoV-2 spike protein-mediated cell growth signaling participates in adverse cardiovascular/pulmonary outcomes seen in COVID-19. This mechanism may provide new therapeutic targets to combat the SARS-CoV-2 infection and COVID-19.

## Abbreviations

ACE2: angiotensin-converting enzyme 2;
ARDS: acute respiratory distress syndrome;
COVID-19: coronavirus disease 2019;
RBD: receptor-binding domain;
SARS-CoV-2: severe acute respiratory syndrome coronavirus 2

## References

[1] Su S, Wong G, Shi W, Liu J, Lai ACK, Zhou J, Liu W, Bi Y, Gao GF. Epidemiology, genetic recombination, and pathogenesis of coronaviruses. Trends Microbiol. 24 (2016) 490–502.

[2] Satija N, Lal SK. The molecular biology of SARS coronavirus. Ann N Y Acad Sci. 1102 (2007) 26–38.

[3] Wu F, Zhao S, Yu B, Chen YM, Wang W, Song ZG, Hu Y, Tao ZW, Tian JH, Pei YY, Yuan ML, Zhang YL, Dai FH, Liu Y, Wang QM, Zheng JJ, Xu L, Holmes EC, Zhang YZ. A new coronavirus associated with human respiratory disease in China. Nature. 579 (2020) 265–269.

[4] Huang C, Wang Y, Li X, Ren L, Zhao J, Hu Y, Zhang L, Fan G, Xu J, Gu X, Cheng Z, Yu T, Xia J, Wei Y, Wu W, Xie X, Yin W, Li H, Liu M, Xiao Y, Gao H, Guo L, Xie J, Wang G, Jiang R, Gao Z, Jin Q, Wang J, Cao B. Clinical features of patients infected with 2019 novel coronavirus in Wuhan, China. Lancet. 395 (2020) 497–506.

[5] Yan R, Zhang Y, Li Y, Xia L, Guo Y, Zhou Q. Structural basis for the recognition of SARS-CoV-2 by full-length human ACE2. Science. 367 (2020) 1444–1448.

[6] Tai W, He L, Zhang X, Pu J, Voronin D, Jiang S, Zhou Y, Du L. Characterization of the receptor-binding domain (RBD) of 2019 novel coronavirus: implication for development of RBD protein as a viral attachment inhibitor and vaccine. Cell Mol Immunol. 17 (2020) 613–620.

[7] Xu Z, Shi L, Wang Y, Zhang J, Huang L, Zhang C, Liu S, Zhao P, Liu H, Zhu L, Tai Y, Bai C, Gao T, Song J, Xia P, Dong J, Zhao J, Wang FS. Pathological findings of COVID-19 associated with acute respiratory distress syndrome. LancetRespirMed. 8 (2020) 420–422.

[8] Wu C, Chen X, Cai Y, Xia J, Zhou X, Xu S, Huang H, Zhang L, Zhou X, Du C, Zhang Y, Song J, Wang S, Chao Y, Yang Z, Xu J, Zhou X, Chen D, Xiong W, Xu L, Zhou F, Jiang J, Bai C, Zheng J, Song Y. Risk factors associated with acute respiratory distress syndrome and death in patients with coronavirus disease 2019 pneumonia in Wuhan, China. JAMA Intern Med. 180 (2020) 1–11.

[9] Li B, Yang J, Zhao F, Zhi L, Wang X, Liu L, Bi Z, Zhao Y. Prevalence and impact of cardiovascular metabolic diseases on COVID-19 in China. Clin Res Cardiol. 109 (2020) 531538.

[10] Yang J, Zheng Y, Gou X, Pu K, Chen Z, Guo Q, Ji R, Wang H, Wang Y, Zhou Y. Prevalence of comorbidities and its effects in patients infected with SARS-CoV-2: A systematic review and meta-analysis. Int J Infect Dis. 94 (2020) 91–95.

[11] Walls AC, Park YJ, Tortorici MA, Wall A, McGuire AT, Veesler D. Structure, function, and antigenicity of the SARS-CoV-2 spike glycoprotein. Cell. 181 (2020) 281–292.e6

[12] Shang J, Wan Y, Luo C, Ye G, Geng Q, Auerbach A, Li F. Cell entry mechanisms of SARS-CoV-2. Proc Natl Acad Sci USA. 117 (2020) 11727–11734.

[13] Wong CM, Cheema AK, Zhang L, Suzuki YJ. Protein carbonylation as a novel mechanism in redox signaling. Circ Res. 102 (2008) 310–318.

[14] Wortzel I, Seger R. The ERK cascade. Distinct functions within various subcellular organelles. Genes Cancer. 2 (2011) 195–209.

[15] Zhang W, Liu HT. MAPK signal pathways in the regulation of cell proliferation in mammalian cells. Cell Res. 12 (2002) 9–18.

[16] Bonjardim CA. Viral exploitation of the MEK/ERK pathway-A tale of vaccinia virus and other viruses. Virology. 507 (2017) 267–275.

[17] Shults NV, Almansour FS, Rybka V, Suzuki DI, Suzuki YJ. Ligand-mediated dephosphorylation signaling for MAP kinase. Cell Signal. 52 (2018) 147–154.

[18] Ding Y, Wang H, Shen H, Li Z, Geng J, Han H, Cai J, Li X, Kang W, Weng D, Lu Y, Wu D, He L, Yao K. The clinical pathology of severe acute respiratory syndrome (SARS): a report from China. J Pathol. 200 (2003) 282–289.

[19] Hwang DM, Chamberlain DW, Poutanen SM, Low DE, Asa SL, Butany J. Pulmonary pathology of severe acute respiratory syndrome in Toronto. Mod Pathol. 18 (2005) 1–10.

